# Measuring CO_2_ assimilation of *Arabidopsis thaliana* whole plants and seedlings

**DOI:** 10.1101/2024.01.22.576682

**Authors:** Ailbhe J. Brazel, Franziska Turck, Diarmuid S. Ó’Maoiléidigh

## Abstract

Photosynthesis is an essential process in plants that synthesizes sugars used for growth and development, highlighting the importance of establishing robust methods to monitor photosynthetic activity. Infrared gas analysis (IRGA) can be used to track photosynthetic rates by measuring the CO_2_ assimilation/release from a plant. Although much progress has been made in the development of IRGA technologies, challenges remain when using this technique on small herbaceous plants such as *Arabidopsis thaliana*. The use of whole plant chambers can overcome the difficulties associated with applying bulky leaf clamps to small delicate leaves, however this introduces the risk of soil-based microorganisms skewing gas exchange measurements. Here, we present a simple method to efficiently perform IRGA on *A. thaliana* plants using a whole plant chamber that removes soil-borne effects from the measurements. We show that this method can be used to detect subtle changes in photosynthetic rates measured at different times of day, under different growth conditions, and between wild-type and plants with deficiencies in the photosynthetic machinery. Furthermore, we show that this method can be used to detect changes in photosynthetic rates even at very young developmental stages such as 10 d-old seedlings. This method contributes to the array of techniques currently used to perform IRGA on *A. thaliana* and can allow for the monitoring of photosynthetic rates of whole plants from young ages.

## Introduction

Plants are autotrophic organisms that use light energy to convert carbon dioxide and water into sugar and oxygen through the process of photosynthesis. Given the importance of photosynthesis to optimal plant growth and development, careful monitoring of photosynthetic rates is desirable. One of the methods used to monitor the rate of photosynthesis is infrared gas analysis (IRGA), which has been used in multiple plant species to track photosynthetic rates in a variety of tissues^1^. Open gas exchange systems are composed of a closed chamber in which a plant organ, or the whole plant, is enclosed while a controlled stream of air passes through it. Infrared sensors are used to measure the CO_2_ levels in the air stream before it enters the chamber and after it leaves the chamber, allowing for the calculation of the rate of CO_2_ assimilation/release by the plant or organ within the chamber.

A number of challenges are presented when applying IRGA to the analysis of photosynthesis of *Arabidopsis thaliana*^2^, a model organism used for molecular genetics approaches. Gas exchange analysis using individual leaf chambers is more easily performed when working with plants with large, expanded leaves or long petioles. *A. thaliana*, however, has relatively small, delicate leaves with short petioles that can be difficult to isolate and are easily damaged by leaf chambers. Furthermore, leaf and plant age influences photosynthetic rates^3^ and it may be difficult to consistently choose comparable leaves, especially when comparing mutant and wild-type plants. To overcome these challenges, whole plant chambers can be used to measure the photosynthetic rates of entire plants as opposed to sections of individual leaves. Importantly, IRGA of entire plants grown on soil with a whole plant chamber will measure both the gas exchange rates that arise from the plant itself and any other organisms, including microorganisms, present in the soil. For this reason, different measures have been adapted to limit the gas exchange from the soil into the chamber including the use of plastic^4^, rubber^5^ or clay^6^ seals. The plant chamber can also be pressurized to cause the air within the chamber to flow out through the soil and thus prevent diffusion of gasses from within the soil into the chamber^7^. However, the release or uptake of CO_2_ by and from the soil sample following these mitigations is not well documented in the literature. Furthermore, manipulation of the plants to place plastic, rubber, or clay seals risks damaging the plant, which can induce changes to the photosynthetic rates^8^ while sample-specific controls for soil-based CO_2_ flux are not described.

There is no standardized plant growth condition for the measurement of CO_2_ assimilation rates, making it difficult to directly compare photosynthetic rates reported for *A. thaliana*^2^. Reported CO_2_ assimilation rates for the ecotype Columbia-0 (Col-0) range from a maximum of ∼4^7^ to ∼20^4^ µmol m⁻² s⁻¹, with many studies reporting a CO_2_ assimilation rate of ∼7 µmol m⁻² s⁻¹^6,9–12^. Measurement of CO_2_ assimilation in *A. thaliana* plants is of interest because of its usefulness as a model plant for molecular genetics. However, to compare between mutant and wild-type plants, specific parameters should be taken into account, such as the length^13–15^ and stability^16^ of the light period in which plants are grown, as well as the time of day at which plants are measured^5,7^, which have been shown to affect CO_2_ assimilation rates assessed by IRGA in plants. For instance, whole *A. thaliana* plants measured continuously over multiple days showed a gradual increase in the rate of CO_2_ assimilation during light periods^5,7^. The effects on photosynthetic rates introduced by the growth regime and time of day could cause challenges when assessing variation in photosynthetic rates between wild-type and mutant plants. Furthermore, many studies employing genome-wide approaches to understanding the topology of the gene regulatory networks (GRNs) underlying photosynthesis use seedling of 10-14 d old plants^17–19^. However, most studies of *A. thaliana* CO_2_ assimilation use plants that are at least 4 weeks old, and often older^4–6,12,20,21^.

Here, we employ a simple and efficient method to remove the effects of contamination from the soil during IRGA in whole *A. thaliana* plants. With this method, we were able to detect differences in CO_2_ assimilation rates between wild-type Col-0 plants measured at different times of day and in different growth conditions. We validate this method by performing IRGA on two mutants with defects in photosynthetic activity. Importantly, we show that growth conditions can influence, and therefore potentially mask or uncover, changes in photosynthetic rates in these mutants. Finally, we employ this method to successfully detect defects in photosynthetic rates in 10 d old *A. thaliana* seedlings, a developmental stage that has not previously been measured with IRGA to our knowledge, but one that is often used to characterize GRNs through genome-wide approaches.

## Results

### Testing a novel “empty pot” method to measure CO_2_ assimilation rates of various genotypes in different growth conditions

To assess the impact of growth conditions on gross photosynthesis for *A. thaliana*, we measured CO_2_ assimilation using IRGA of plants grown in three different daylengths including continuous light (CL), long day (LD) and short day (SD) conditions (Fig. 1A, Table S1). We performed measurements of whole plants in each daylength in the morning (Early) and in the afternoon (Late) (Fig. 1A; see Materials and Methods) using a light-curve program (0-1000 µmol m⁻² s⁻¹ in 250 µmol m⁻² s⁻¹ increments) in a small plant chamber. For reasons outlined in the introduction, we employed a novel “empty pot” (EP) method in which we obtained CO_2_ assimilation rates by subtracting the IRGA values for the soil alone from the measurements of the whole plant and the soil (Fig. 1B; see Materials and Methods). We used several genotypes including Col-0 “wild-type” controls, a mutant for *CHLOROPLAST IMPORT APPARATUS 2* (*CIA2*), whose product controls the expression of genes involved in protein import to, and translation in, the chloroplast^19,22^, and a double mutant of *GOLDEN-2-LIKE 1* (*GLK1*) and *GLK2*, whose products are involved in chloroplast development^23^.

**Fig. 1.**
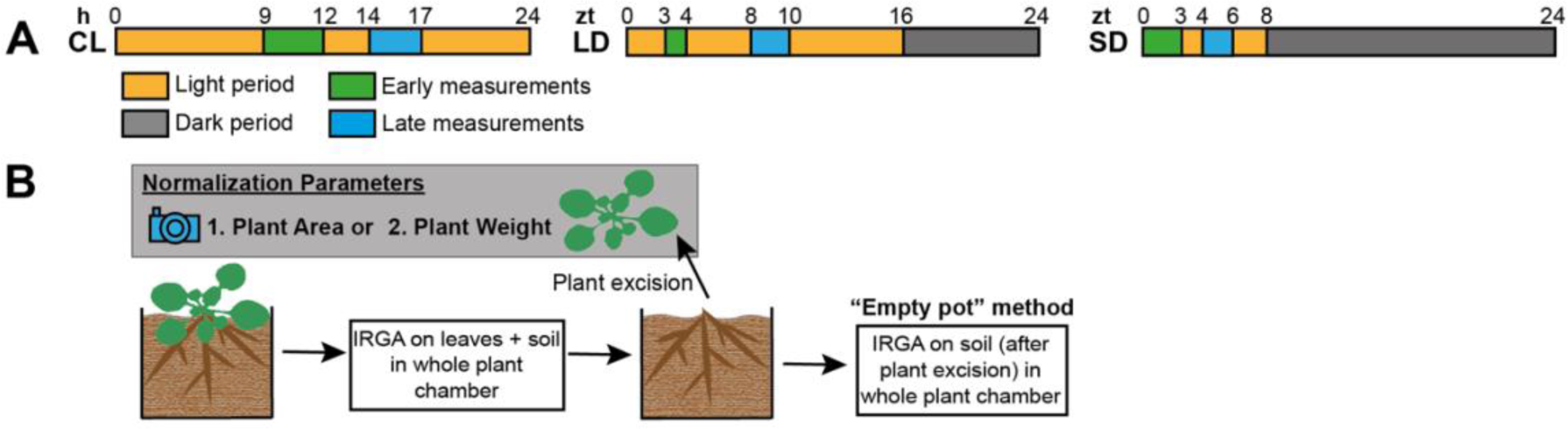
Strategy for measuring photosynthetic rates by IRGA. (A) *A. thaliana* plants were grown in three growth conditions with different light regimes. The light regimes, “continuous light” (CL), “long days” (LD) and “short days” (SD) are indicated by orange light period and gray dark period. Within each light regime, CO_2_ gas exchange measurements were started at “Early” (green) or “Late” (blue) times of day, as indicated in the figure. See Materials and Methods for more details. (B) CO_2_ gas exchange was measured by IRGA of whole pots containing an *Arabidopsis* plant and the soil in which it was grown. The aerial section of the plant was then excised and photographed to measure its area and the fresh weight. For the EP method, IRGA was then performed on the pot without the aerial section of the plant. CO_2_ gas exchange data was normalized using either plant area or plant fresh weight measurements. The CO_2_ gas exchange data from the pot without the aerial section of the plant was subtracted from the data from the pot containing the aerial section to obtain measurements of leaf photosynthesis that removed the contribution of respiration originating from the soil.

We initially used an area-based method (Fig. 1B) to normalize the CO_2_ assimilation rates where we measured the area of the plant before IRGA (see Materials and Methods). To test if CO2 gas exchange occurs in the soil we first analyzed IRGA data from the “empty pots”, where we excised the aerial section of the plant before performing gas exchange measurements (Fig. S1). CO_2_ assimilation rates were negative for all genotypes tested indicating a strong source of respiration from the “empty pots” (e.g. microorganisms) (Fig. 1B, Fig. S1). There was a high degree of variation for measurements within a single growth condition and genotype suggesting that the respiration levels are unique to each pot (Fig. S1). Notably, the “empty pot” respiration was different between genotypes with Col-0 ranging from −6.7 to −3.9 µmol m⁻² s⁻¹, *cia2-1* ranging from −8.8 to −4.5 µmol m⁻² s⁻¹, and *glk11-1 glk2-1* ranging from −17.2 to −9 µmol m⁻² s⁻¹ (Fig. S1, Table S2). Therefore, the respiration originating from the soil is unique to each genotype and significantly influences respiration rates depending on the growth condition.

In each growth condition, *glk1-1 glk2-1* adult plants were the smallest in size followed by *cia2-1*, suggesting a reduction in photosynthetic capacity, in keeping with the pale phenotypes of *cia2-1* and *glk1-1 glk2-1*, or an increase in respiration (Fig. 2A). Using the EP method combined with an area-based normalization there was a clear difference in CO_2_ assimilation rates (Δ1.5 – 2.2 µmol m⁻² s⁻¹) between the mutant genotypes and Col-0 under CL condition at all light intensities (*p.adj* < 0.03; Fig. 2B, Table S2). Under LD conditions, CO_2_ assimilation of *cia2-1* was lower than Col-0 (Δ1.2 – 1.9 µmol m⁻² s⁻¹, *p.adj* < 0.04) at light intensities from 500 – 1000 µmol m⁻² s⁻¹ with *glk1-1 glk2-1* CO_2_ assimilation only being reproducibly lower than Col-0 (Δ1.2 µmol m⁻² s⁻¹, *p.adj* = 0.04) at the 250 µmol m⁻² s⁻¹ light intensity (Fig. 2C, Table S2). The largest magnitude of CO_2_ assimilation rate differences was between Col-0 and the two mutant genotypes under SD conditions (Δ1.9 – 3.1 µmol m⁻² s⁻¹; *p.adj* < 0.05; Fig. 2D, Table S2). This demonstrates that CO_2_ assimilation rates are influenced by interactions between growth conditions and genotype.

**Fig. 2.**
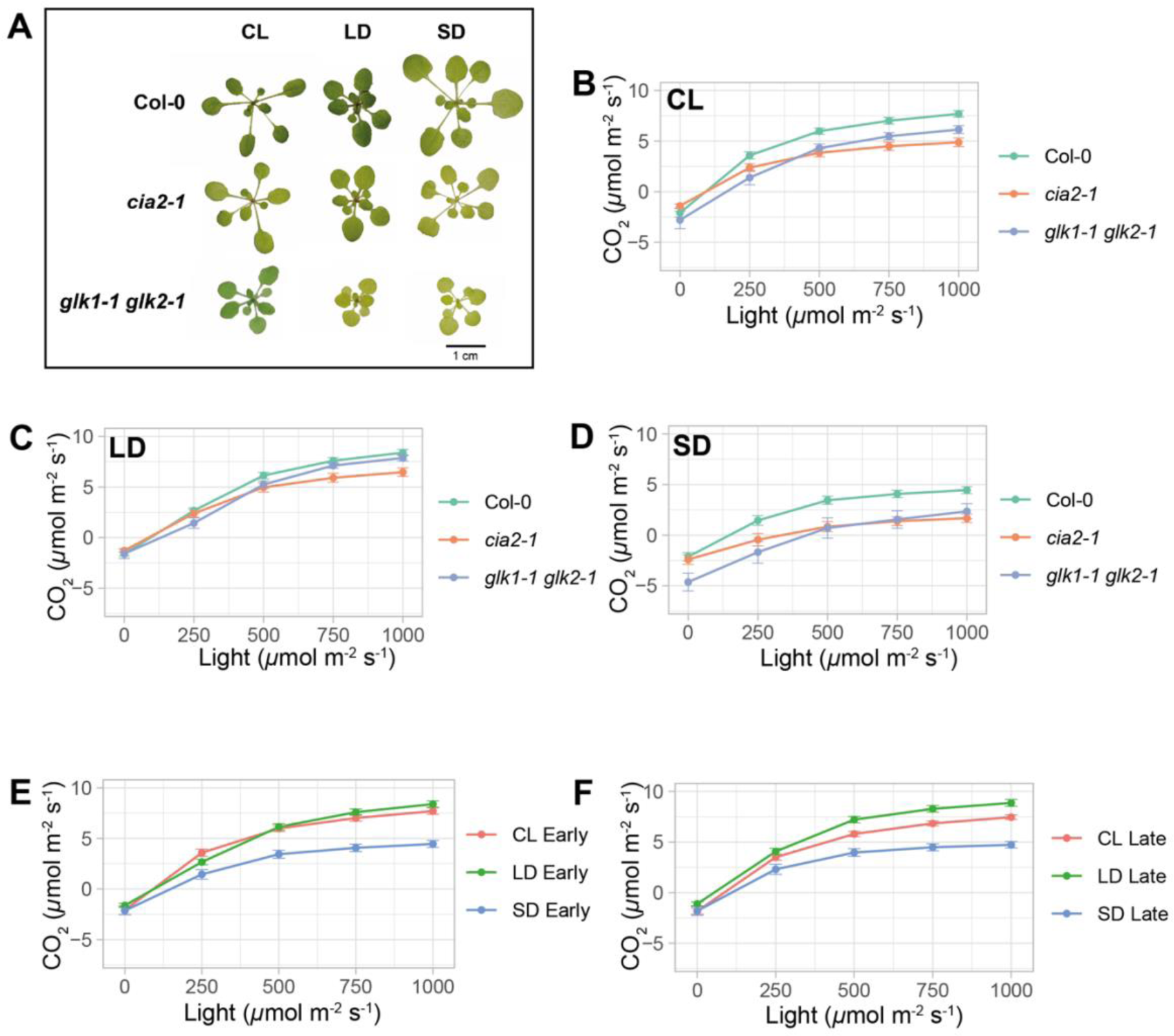
Gas exchange measurements in genotypes with defects in photosynthesis. (A) Representative photographs of Col-0, *cia2-1* and *glk1-1 glk2-1* adult plants grown in CL and LD conditions for 3 w, and SD conditions for 3.5 w. (B-D) CO_2_ gas exchange rates measured using the EP method at “Early” time points at varying light intensities are shown for whole adult plants grown in (B) CL, (C) LD and (D) SD conditions. (E-F) CO_2_ gas exchange rates at varying light intensities are shown for whole Col-0 plants grown in continuous light (CL), long day (LD) and short day (SD) conditions measured at (E) “Early” and (F) “Late” time points using the EP method. Error bars in are s.e.m. of (B-D) 6-18 individual plants from 3-9 biologically independent replicates and (E-F) 12-18 individual plants from 6-9 biologically independent replicates. See Table S2 for statistical analysis of the data.

Though the area based normalization worked well with the EP method, we also tested whether a fresh weight-based measurement would also be suitable (Fig. 1B). In comparison to the area-based assessment of mean CO_2_ assimilation rates (Fig. 2B), the differences between Col-0 and *cia2-1* were compressed (Δ0.6 – 1.2 µmol m⁻² s⁻¹, *p.adj* = 0.06 – 0.51; Fig. S2A, Table S2). In contrast, *glk1-1 glk2-1* CO_2_ assimilation rates were reproducibly higher than Col-0 at the two highest light intensities (Δ1.6 – 1.9 µmol m⁻² s⁻¹, *p.adj* < 0.021; Fig. S2A, Table S2). The EP method reduced the variation for each measurement in comparison to when the EP method was not used (Fig. S2B-C). We concluded that the area-based normalization was more suitable as fresh-weight measurements introduce variation in due to moisture content, which could vary between genotypes. A dry-weight normalization is an alternative possibility, however, since the area-based normalization produced robust results, we continued with this approach.

We also tested another recommended method to account for gas exchange from the soil by using pressure differentials. Here, we applied a 50% leak to create pressure in the small plant chamber and prevent gasses diffusing from the soil into the chamber. With this method, differences in dark respiration of each genotype were larger when compared to the EP method (Fig. S2D). Mean CO_2_ assimilation rates were lower for *glk1-1 glk2-1* compared to Col-0 (*p.adj* > 0.1, Δ4.8 – 5.3 µmol m⁻² s⁻¹) in all light conditions whereas mean CO_2_ assimilation rates for *cia2-1* was higher compared to Col-0 (*p.adj* > 0.64, Δ1.3 – 2.6 µmol m⁻² s⁻¹). Notably, the variation of each measurement was much higher compared to the EP method meaning that the differences between the means of the mutant the genotypes relative to Col-0 were not supported by statistical analysis (Table S2). Furthermore, CO_2_ assimilation rates of Col-0 measured with the pressure method were consistently lower than values measured with the EP method (*p.adj* < 0.06, Δ1.2 – 4.3 µmol m⁻² s⁻¹; Fig S2E-F, Table S2), likely due to a loss of gasses due to the leak that was applied.

### Testing CO_2_ assimilation at different times of day

Differences in CO_2_ assimilation rates were also observed between plants measured at “Early” and “Late” times of day. Under CL and LD conditions, Col-0 plants that were measured at “Early” had very similar CO_2_ assimilation rates in all light intensities (*p.adj* > 0.1, Δ0.1 – 0.9 µmol m⁻² s⁻¹; Fig. 2E, Table S2). However, CO_2_ assimilation of LD grown plants measured at “Early” timepoints were lower than LD grown plants measured at “Late” timepoints in light intensities from 500 – 1000 µmol m⁻² s⁻¹ (*p.adj* < 0.004, Δ1.4 µmol m⁻² s⁻¹; Fig. 2F, Table S2). SD grown plants had consistently lower CO_2_ assimilation rates compared to both CL and LD grown plants regardless of the time of day the plants were measured (*p.adj* < 0.04, Δ1.2 – 4.1 µmol m⁻² s⁻¹; Fig. 2E-F, Fig. S2G, Table S2). The differences of CO_2_ assimilation rates between growth conditions was further verified in all light intensities using an individual leaf clamp (*p.adj* < 0.01, Δ1.2 – 2.6 µmol m⁻² s⁻¹; Fig. S2H, Table S2).

### Testing methods to detect CO_2_ assimilation rates in *A. thaliana* seedlings

To our knowledge, CO_2_ flux experiments have not yet been performed on *A. thaliana* seedlings, which is desirable as 10-14 d old seedlings are widely used in biochemical and genome-wide approaches to study protein activity and GRNs. To test whether differences in CO_2_ assimilation rates could be detected in seedlings with defects in chloroplast development, we performed IRGA on 10 d old Col-0, *cia2-1* and *glk1-1 glk2-1* seedlings grown under CL growth conditions on both soil-filled pots and MS-agar plates (see Materials and Methods). As with the adult plants (Fig. 2A), both the *cia2-1* and *glk1-1 glk2-1* seedlings were paler in appearance than Col-0 (Fig. 3A). Remarkably, despite the small size of the 10 d old *A. thaliana* seedlings (Fig. S3), we were able to detect differences in CO_2_ gas exchange rates in plants grown on soil in pots (Fig. 3B). The rate of CO_2_ assimilation in *cia2-1* seedlings was lower than that of Col-0 seedlings in light intensities from 500 – 1000 µmol m⁻² s⁻¹ (*p.adj* < 0.0001, Δ1.4 – 1.6 µmol m⁻² s⁻¹; Fig. 3B, Table S2). Interestingly, there was less dark respiration observed of *glk1-1 glk2-1* seedlings (−1.2 µmol m⁻² s⁻¹, respectively) compared to Col-0 (−2.3 µmol m⁻² s⁻¹, *p.adj* < 0.0001) (Fig. 3B, Table S2). We were, however, unable to detect differences in CO_2_ gas exchange rates in plants grown on plates (Fig. 3C, Table S2), possibly due to the smaller average size of seedlings grown on plates compared to pots (Fig. S3).

**Fig. 3.**
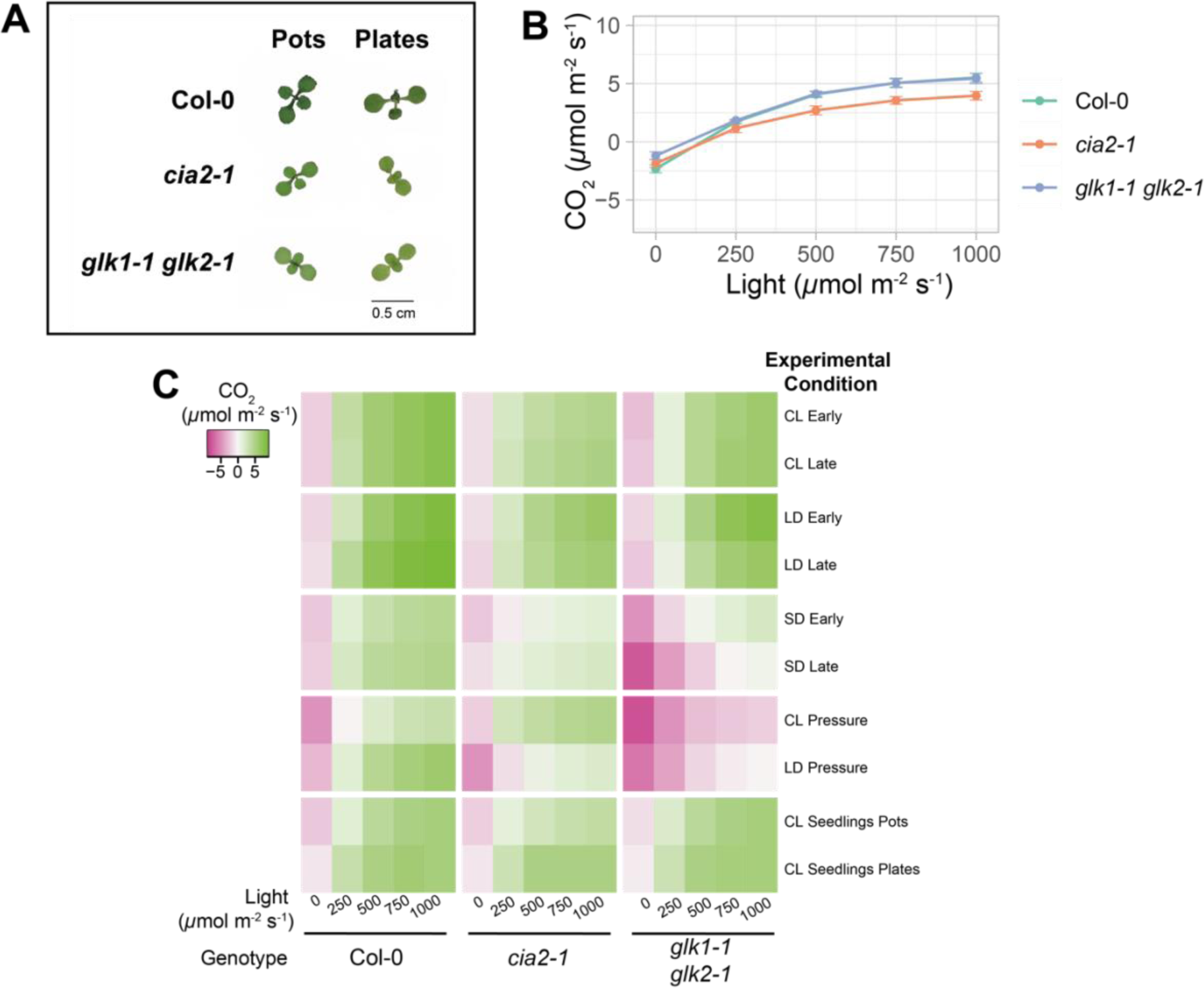
Comparison of gas exchange measurements across genotypes, plant age, growth conditions and measurement time. (A) Representative photographs of Col-0, *cia2-1* and *glk1-1 glk2-1* adult plants grown in CL conditions for 10 d in pots/plates. (B) CO_2_ gas exchange rates at varying light intensities are shown for 10 d seedlings grown in CL conditions in pots. Error bars are s.e.m. of 11-14 individual pots, each containing 5-16 seedlings, from 4 biologically independent replicates. (C) Heatmap summary of average CO_2_ gas exchange rates at varying light intensities for Col-0, *cia2-1* and *glk1-1 glk2-1* in all experimental conditions tested in this study. See Table S2 for statistical analysis of the data.

## Discussion

Here, we present a simple and useful method for measuring photosynthetic rates using IRGA on whole *A. thaliana* plants. Although leaf clamps can be used to measure CO_2_ assimilation rates, the small leaf size and close rosette structure of *A. thaliana* plants make the use of bulky leaf clamps difficult. If measuring attached leaves, it is therefore necessary to wait until the plants have advanced in age and the leaves have expanded sufficiently before the use of a leaf clamp is possible. The consistent selection of leaves at the same developmental stage is critical as leaves at different ages have different photosynthetic rates^3^. However, this becomes more difficult with older *A. thaliana* plants, particularly when dealing with mutant lines where developmental progression is affected, and leaf size may be reduced (Fig. 2A). It is also important to note that many transcriptomic studies in *A. thaliana* are performed on young plants (10-14 d)^17–19^ and direct comparison of these data with physiological measurements is valuable.

Whole plant chambers have been utilized to overcome the challenges associated with leaf-clamp measurements in *A. thaliana* plants. CO_2_ analysis using whole plant chambers detects respiration/photosynthesis from both the plant of interest and any microorganisms present in the soil unless methods are used to block diffusion from the soil. However, the effectiveness of these methods to prevent contamination from soil-based microorganisms (Fig. S1) is not well described, while some of the methods used are cumbersome, risk damaging the plant tissue, and controls to verify the absence of soil-based contamination of CO_2_ assimilation rates are not easily included.

The method we present here is a simple way to ensure contamination from soil-based microorganisms are removed from CO_2_ assimilation analysis (Fig. 1). By subtracting values measured from an “empty pot” from values obtained from a pot containing the plant of interest, we completely remove the contribution of soil-based microorganisms from our photosynthetic rate analysis. The presented method produced estimates of gross photosynthetic rates similar to those produced with a leaf clamp (Fig. S2H) and to those previously reported^6,9–12^.

We present a large-scale dataset in which we use this method to measure CO_2_ assimilation rates in wild-type and photosynthesis mutant adult plants measured at different times of day in different growth conditions, as well as 10 d old seedlings (Fig. 3C, Table S1). This method improves the reproducibility of gas exchange analysis and reduces the rate of error in these measurements associated with contamination from soil-based microorganisms (Fig. 2, Fig. S2). We show that this method is sensitive enough to detect changes in photosynthetic rates caused by differing growth conditions, time of day and mutation of photosynthetic machinery (Fig. 2, Fig S2). Finally, we use this method to measure photosynthetic rates in 10 d old seedlings and show that *cia2-1* mutants also display a reduced CO_2_ assimilation rate at this young developmental stage (Fig. 3B). The reproducible results obtained with this method can help improve detection of defects in photosynthesis at multiple developmental stages and highlight the importance of consistent conditions when carrying out these measurements.

## Materials and methods

### Plant growth and materials

Seeds were sown on autoclaved (60 min at 125°C) potting medium (10:6:4, compost:vermiculite:perlite) in 6 cm pots, or on 3.5 cm plates with 0.5x MS agar. Seeds were stratified for at least two days in the dark at 4°C. Plants were then moved to growth rooms in continuous light (CL), long day (LD) 16:8 h light:dark, short day (SD) 8:16 h light:dark or neutral day (ND) 12:12 h light:dark conditions, as described in the text and Table S1. The specific ages that plants were measured at is described in Table S1. For adult plant measurements, 1 plant was grown per pot while for 10 d seedling measurements 5-17 seedlings were planted per pot or plate. The plant lines used in this study are *Arabidopsis thaliana* ecotype Col-0, *cia2-1*^22^ (N6522) and *glk1-1 glk2-1*^23^ (N9807). These plant lines were acquired from the Nottingham Arabidopsis Stock Centre.

### Infrared gas exchange analysis

Plants were grown to the age indicated in Table S1 and well-watered before their photosynthetic CO_2_ assimilation rates were measured using an LI-6800 Portable Photosynthesis System (LI-COR). For plants grown in CL growth conditions, “Early” measurements were initiated between approximately 09:00 and 12:00 while “Late” measurements were initiated between 14:00 and 17:00. For plants grown in LD growth conditions, “Early” measurements were initiated between approximately Zeitgeber Time (ZT) 3 and ZT4, while “Late” measurements were initiated between ZT8 and ZT10. For plants grown in SD growth conditions, “Early” measurements were initiated between approximately ZT0 and ZT3, while “Late” measurements were initiated between ZT4 and ZT6.

CO_2_ assimilation measurements were carried out using a small plant chamber with a mounted large light source. Where indicated, a leaf clamp chamber with a 6 cm^2^ aperture was used for IRGA. Plants were placed into the sealed small plant chamber with the light source switched off and allowed to acclimate for at least 5 min in the dark, or until the CO_2_ sample readings stabilized. A light response curve was performed using the following settings; a flow rate of 600 µmol s⁻¹, sample water vapor set point of 19 mmol mol⁻¹, reference CO_2_ of 400 µmol⁻¹, mixing fan speed of 10,000 rpm, heat exchanger temperature setpoint of 22°C, chamber pressure 0.2 kPa, light source composition 90% red 10% blue light, light intensity curve set points 0, 250, 500, 750 and 1000 µmol m⁻^2^ s⁻¹ with a 120 – 180 s wait time between set points (no early match allowed). For experiments with the individual leaf clamp chamber, plants were grown to 3-4 weeks old and the same light response curve was applied with the following settings on the two largest most expanded leaves per plant; a flow rate of 400 µmol s⁻¹, relative humidity of air setpoint of 55-60%, reference CO_2_ of 400 µmol⁻¹, mixing fan speed of 10,000 rpm, heat exchanger temperature setpoint of 20°C, chamber pressure 0.2 kPa and light source composition 90% red 10% blue light. When the “Pressure” method was used, a leak of approximately 50% or 300 µmol m⁻^1^ was applied to release valve beneath the pot. When the “empty pot” (EP) method was used, the aerial part of the plant was excised from the pot without disturbing the soil and without removing the pot from the chamber after the light response curve measurement was completed. The chamber was then sealed again with the “empty pot” within and another light response curve measurement was performed.

### Measurement of Plant Size

Following the measurement of photosynthetic rate, the plant was imaged and weighed. Plant images were processed in Adobe Photoshop 2022. The plant area was measured by using the magic wand and magnetic lasso tool to quickly select the plant tissue. For the seedling area measurements, average seedling size was calculated by dividing the total area calculated for all the seedlings in an individual pot/plate by the number of seedlings measured.

### Data analysis

CO_2_ assimilation rates were normalized on an area (µmol CO_2_ m⁻² s⁻¹) or fresh-weight (µmol CO_2_ mg⁻^1^ s⁻¹) basis, as described in the text. When the EP method was used, the CO_2_ assimilation rate measured for the “empty pot” at each light intensity was subtracted from the value obtained for the pot containing the plant at each corresponding light intensity.

Data were analyzed using Prism 10 for macOS, Version 10.1.0 (264), graphs were made using the *ggplot2* package from R studio software^24^ and figures were edited for appearance using Adobe Illustrator 2022. For multiple comparisons, data were analyzed using a two-way analysis of variance (ANOVA). Post-hoc tests were employed if a significant result was obtained from the above tests (p < 0.05). Following an ANOVA test, the Šídák multiple comparisons test was used when two groups were being compared, while the Tukey’s multiple comparisons test was used when three groups were being compared. The heatmap was generated using the Heatmapper web tool^25^.

## Acknowledgements

We thank Dr. Emmanuelle Graciet for use of reagents and equipment to perform experiments.

---

**Supplementary Fig. 1.**
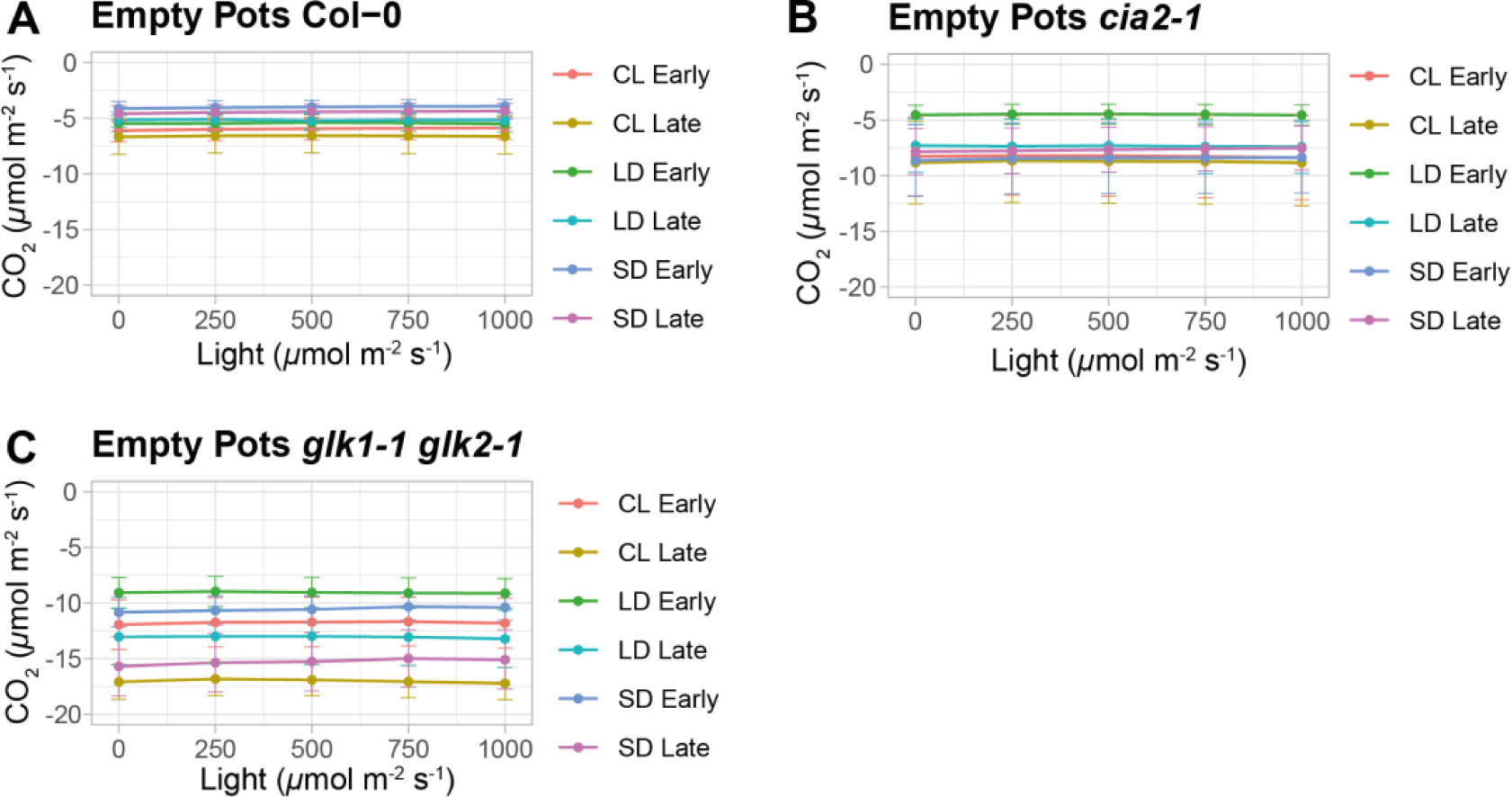
CO_2_ assimilation rate measurements of “empty pots” following excision of aerial section of plants from pots. (A-C) *A. thaliana* plants were grown in CL, LD or SD growth conditions and measured at “Early” or “Late” timepoints, as described in Fig. 1. CO_2_ assimilation rates were measured at different light intensities for “empty pots” i.e. pots which still contained soil from which the aerial section of the plant had freshly been removed. Measurements of “empty pots” were made following excision of (A) Col-0, (B) *cia2-1* and (C) *glk1-1 glk2-1*. Error bars in are s.e.m. of 6-18 plants from 3-9 biologically independent replicates. See Table S2 for statistical analysis of the data.

**Supplementary Fig. 2.**
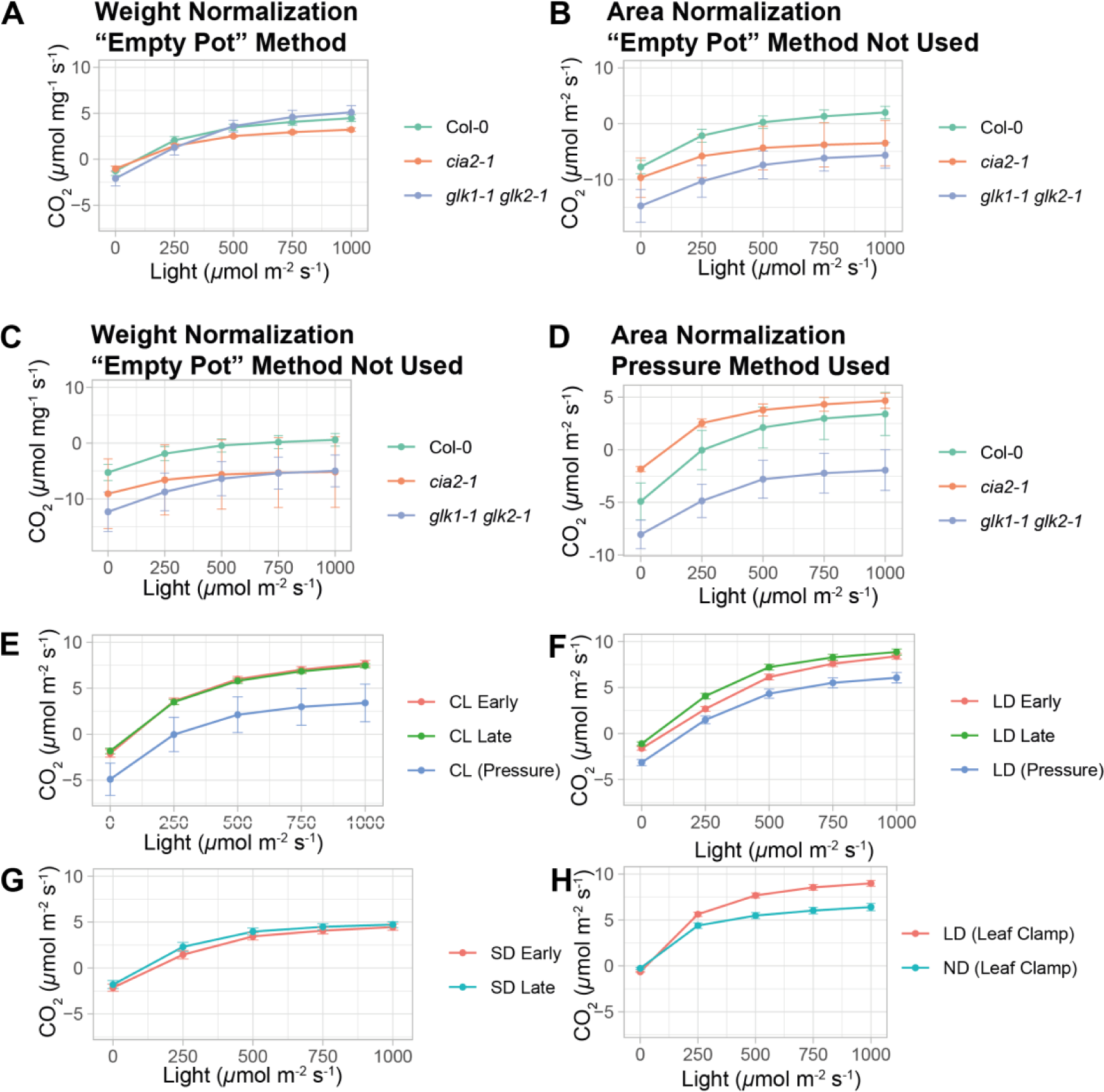
Analysis methods for CO_2_ assimilation rate. (A-D) CO_2_ assimilation rates at varying light intensities are shown for whole adult plants grown in CL conditions and measured at “Early” timepoints. Photosynthetic rates were normalized (A) on a weight basis using the EP method, (B) on an area basis without using the EP method, (C) on a weight basis without using the EP method and (D) on an area basis using the “pressure” method. (E-G) CO_2_ assimilation rates at varying light intensities are shown for whole Col-0 plants measured at “Early” and “Late” timepoints, or with the “pressure” method, grown in (E) CL, (F) LD and (G) SD conditions. (H) CO_2_ gas exchange rates at varying light intensities are shown for individual leaves from Col-0 plants grown in LD and neutral day (ND) conditions measured using a leaf clamp. Error bars are s.e.m. of (A-D) 6-14 individual plants from 3-7 biologically independent replicates, (E-G) 12-18 individual plants from 6-9 biologically independent replicates and (H) 17-22 leaves from 4 biologically independent replicates. See Table S2 for statistical analysis of the data.

**Supplementary Fig. 3.**
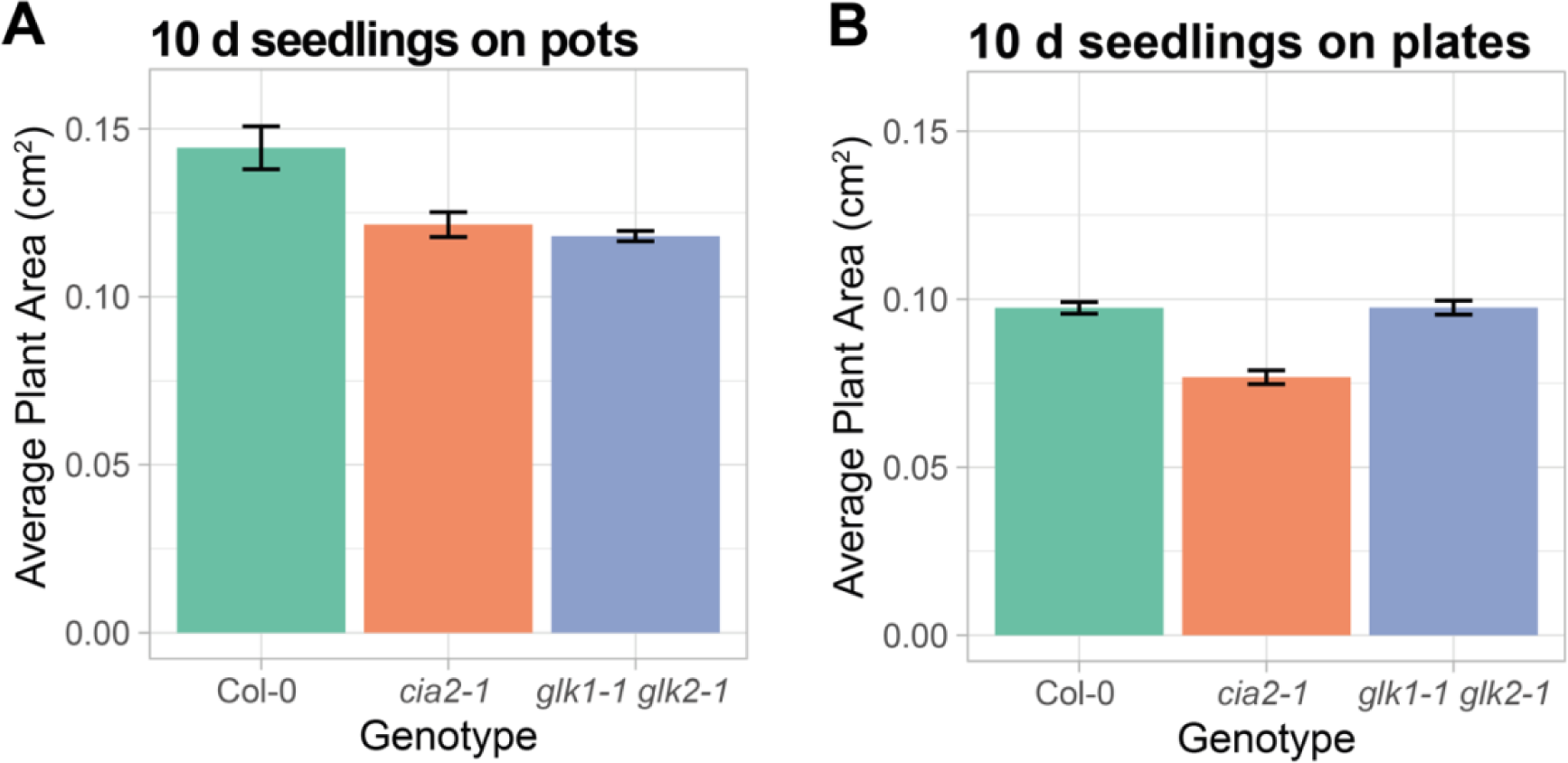
Average plant areas of 10 d seedlings grown in CL conditions. (A) Average 10 d old seedlings plant sizes are shown for 128, 116 and 138 seedlings for Col-0, *cia2-1* and *glk1-1 glk2-1* plants, respectively, grown in pots, over 3-4 independent biological replicates. (B) Average 10 d old seedlings plant sizes are shown for 58, 57 and 64 seedlings for Col-0, *cia2-1* and *glk1-1 glk2-1* plants, respectively, grown on plates, over 1-2 independent biological replicates. Error bars represent s.e.m.

